# Ecophysiology and genomics of the brackish water adapted SAR11 subclade IIIa

**DOI:** 10.1101/2022.08.02.502558

**Authors:** V. Celeste Lanclos, Anna N. Rasmussen, Conner Y. Kojima, Chuankai Cheng, Michael W. Henson, Brant C. Faircloth, Christopher A. Francis, J. Cameron Thrash

**Author notes:** Correspondence: J. Cameron Thrash, University of Southern California, Department of Biological Sciences, 3616 Trousdale Pkwy AHF 107 Los Angeles, CA 90089.

## Abstract

The Order Pelagibacterales (SAR11) is the most abundant group of heterotrophic bacterioplankton in global oceans and comprises multiple subclades with unique spatiotemporal distributions. Subclade IIIa is the primary SAR11 group in brackish waters and shares a common ancestor with the dominant freshwater IIIb (LD12) subclade. Despite its dominance in brackish environments, subclade IIIa lacks systematic genomic or ecological studies. Here, we combine closed genomes from new IIIa isolates, new IIIa MAGS from San Francisco Bay (SFB), and 466 high-quality publicly available SAR11 genomes for the most comprehensive pangenomic study of subclade IIIa to date. Subclade IIIa represents a taxonomic family containing three genera (denoted as subgroups IIIa.1, IIIa.2, and IIIa.3) that had distinct ecological distributions related to salinity. The expansion of taxon selection within subclade IIIa also established previously noted metabolic differentiation in subclade IIIa compared to other SAR11 subclades such as glycine/serine prototrophy, mosaic glyoxylate shunt presence, and polyhydroxyalkanoate synthesis potential. Our analysis further shows metabolic flexibility among subgroups within IIIa. Additionally, we find that subclade IIIa.3 bridges the marine and freshwater clades based on its potential for compatible solute transport, iron utilization, and bicarbonate management potential. Pure culture experimentation validated differential salinity ranges in IIIa.1 and IIIa.3 and provided the first IIIa cell size and volume data. This study is an important step forward for understanding the genomic, ecological, and physiological differentiation of subclade IIIa and the overall evolutionary history of SAR11.

## Introduction

The SAR11 clade (Pelagibacterales) is a diverse order of bacterioplankton that constitutes up to 40% of heterotrophic bacteria in surface global oceans (1, 2). The clade encompasses multiple subclades that exhibit unique spatiotemporal distributions in global waters corresponding to the group’s phylogenetic structure (1, 3). Much of what is known about SAR11 comes from subclade I and the well-characterized strains HTCC1062 and HTCC7211 (4–6). Studies focused on these organisms and other genomes within Ia defined SAR11 as canonical genome-streamlined oligotrophic marine heterotrophs (7–9) with specific nutrient requirements (10), simple regulatory systems (7, 11, 12), auxotrophies for key amino acids and vitamins (13, 14), partitioning of carbon flow for assimilation or energy based on external nutrient concentrations (15), and sensitivity to purifying selection within closely related populations (16). Studies of non-Ia SAR11 subclades have provided evidence of additional subclade-specific genomic adaptations and biogeography. For example, subclade Ic contains subtle genomic changes such as amino acid composition, increased intergenic spacer size, and genes encoding for cell wall components as likely adaptations to the bathypelagic (17). Some subclade II members possessed genes for nitrate reduction in oxygen minimum zones, providing the first evidence of facultative anaerobic metabolism in SAR11 (18). The freshwater LD12/IIIb subclade was recently cultivated and its growth in low brackish salinities and loss of osmoregulation genes provides a hypothesis for SAR11 adaptation into freshwater ecosystems (19, 20).

Another important SAR11 subclade, IIIa, which shares a most recent common ancestor with the freshwater LD12/IIIb group (3, 19) (hereafter LD12), has received comparatively little attention despite being a key group to study the evolutionary transition of SAR11 from marine to fresh water. To date, there are only two reported isolates, HIMB114 (8) and IMCC9063 (21) but this lack of systematic study is not indicative of IIIa’s relevance in global aquatic systems. IIIa is the most abundant SAR11 subclade in brackish waters and its distribution varies based on salinity and phylogenetic position, with two primary branches represented by the two isolates and their genomes (22, 23). In a survey of the Baltic Sea, the IMCC9063-type of SAR11 was the more abundant representative in brackish waters (salinity < 10) while the HIMB114-type peaked in high-brackish to marine salinities (22). A similar trend has also been seen across northern Gulf of Mexico estuaries in which multiple operational taxonomic units (OTUs) of SAR11 IIIa were separated ecologically by salinities above and below ∼10 (23). In the San Francisco Bay (SFB), a 16S rRNA amplicon OTU-based study also found subclade IIIa to dominate at mesohaline salinities (24). Additionally, the two established branches of IIIa were separated by temperature and latitude in polar versus temperate waters (25). Despite evidence of niche separation based on their environmental distributions, the temperature and salinity tolerances of these organisms have not be tested experimentally.

There is a comparative paucity of information about subclade IIIa relative to other SAR11, and only limited information has been gleaned from studies using comparative genomics thus far. Neither IIIa representative contains a complete glycolytic pathway, though the neighboring subclade LD12 contains a typical EMP pathway (19) and some subclade I representatives have a variant of the ED pathway (26). While all SAR11 members are reliant on reduced sulfur, neither HIMB114 nor IMCC9063 have the genomic potential to use DMSO or DMSP like other SAR11 strains (15, 27–29). The extensive C1 metabolism found in other SAR11 strains is also lacking in IIIa genomes (17). Contrary to other SAR11 members, HIMB114 and IMCC9063 have been reported to contain *ser*ABC for glycine/serine prototrophy and IMCC9063 also contains a *tenA* homolog not found in subclade I that may allow for AmMP rather than HMP to serve as a thiamin source (14). Together, these genomic predictions suggest that IIIa is fundamentally different from other SAR11 clades in some aspects of metabolic potential which aligns with the general SAR11 trend of phylogeny reflecting the unique ecology and genomic novelty of particular clades. Furthermore, 16S rRNA gene and phylogenomic trees indicate at least three separate IIIa subgroups instead of only two, raising questions about possible additional genomic and ecological diversification within IIIa (3, 30).

To improve our understanding of the genomic, ecological, and physiological variation present in SAR11 subclade IIIa, we conducted a comprehensive study leveraging new isolates, three closed genomes from these strains, and an additional 468 SAR11 genomes that included new and publicly available metagenome-assembled genomes (MAGs), single-amplified genomes (SAGs), and 1059 metagenomic samples from a variety of aquatic habitats. We examined the pangenomics and global ecology of the group as well as pure culture physiology from two of our isolates. Our results provide strong evidence for three genera within IIIa (IIIa.1, IIIa.2, and IIIa.3) whose ecological distribution is defined at least partially by salinity. We define the genomic adaptations that separate IIIa from the rest of SAR11, the three subgroups within IIIa from each other, and partially characterize the physiology and morphology of two isolates from the IIIa branches with cultured representatives. Our SAR11 IIIa strains grown in defined and complex artificial seawater medium, as well as their genomes, provide new opportunities for detailed study of this group.

## Materials and Methods

### Isolation, genome sequencing, and assembly

All strains were isolated using high throughput dilution-to-extinction methods and identified through 16S rRNA gene sequences as previously reported (25, 31). DNA for strain LSUCC0261 was sequenced using Illumina HiSeq after library preparation as previously reported (19) at the Oklahoma Medical Research Facility. DNA for strains LSUCC0664 and LSUCC0723 was sent to the Argonne National Laboratory Environmental Sample Preparation and Sequencing Facility for library preparation and sequencing. We trimmed reads with Trimmomatic v0.36 and assembled trimmed reads for all genomes with SPAdes v3.10.1 (32) using default parameters with coverage cutoff set to “auto”. We verified closure of the genomes and checked the assemblies for contamination using CheckM v1.0.5 (33) with “lineage_wf”. See **Supplemental Text** for detailed methods on isolation, sequencing, assembly, binning, and genome closure verification.

### Comparative genomics, and genome characteristics

Subgroups within SAR11 were delineated using phylogenetic branching (**Supplemental Text**), 16S rRNA gene BLAST identity, and average and average amino acid identity (AAI) (https://github.com/dparks1134/CompareM, default settings). Comparative genomics was completed using Anvi’o version 7.1 (34, 35) with the pangenomics workflow (https://merenlab.org/2016/11/08/pangenomics-v2) as previously reported (36). We also searched for bacteriophage in the assembled genomes of LSUCC0261, LSUCC0664, and LSUCC0723 using the Virsorter ‘Virome’ and ‘RefSeq’ databases (37). Lastly, we used CheckM v1.0.5 (33) output values for genome characteristics (coding density, GC%, predicted genes, and estimated genome size) comparison. We estimated the genome size of non-closed genomes that were at least 80% complete by multiplying the number of base pairs in the genome assembly by the inverse of the estimated completion percentage (**Table S1**).

### Competitive metagenomic read recruitment

To examine the distribution of genomes in aquatic systems, we selected 1,059 metagenomes for read recruitment from the following regions: Baltic Sea, Chesapeake Bay, Columbia River, Black Sea, Gulf of Mexico, Pearl River, Sappelo Island, San Francisco Bay, BioGeoTraces, Tara Oceans, and HOT (accession numbers available in **Table S1**). We conducted read mapping and calculation of normalized abundances via Reads Per Kilobase (of genome) per Million (of recruited read base pairs) (RPKM) using RRAP(38).

### Growth experiments

To test the salinity and temperature ranges of our isolates, we grew pure cultures in their isolation medium across a range of ionic strengths and temperatures in the dark without shaking. To test for various C, N, and S substrates that could be used by LSUCC0261, we grew the culture in a modified JW2 medium that contained a single carbon, nitrogen, and sulfur source(**Table S1)** in 96 × 2.1 mL well PTFE plates (Radleys, Essex, UK). Concentrations for the nutrient sources were added to mimic those in the original minimal media as follows: carbon 500 nM, nitrogen 5 µM, sulfur 90 nM for cysteine and methionine and 500 nM for taurine. After three sequential transfers of the plates every 3-4 weeks, we transferred any wells that showed a cell signature on the flow cytometer to flasks in triplicate with the corresponding C/N/S mixtures and a higher concentration of the carbon substate (50 µM). All cultures were re-checked for purity after the experiment concluded via Sanger sequencing of the 16S rRNA gene as described (31). Cell concentrations were enumerated using a Guava EasyCyte 5HT flow cytometer (Millipore, Massachusetts,USA) with previously reported settings (19, 31). Growth rates were calculated using sparse-growth-curve (41).

### Electron microscopy and cell size estimates

LSUCC0261 was grown to 10^6^ cells mL^-1^ and 50mL of culture was fixed with 3% glutaraldehyde at 4’C overnight. Cells were filtered onto a 0.2µm Isopore polycarbonate membrane filter (MilliporeSigma) and dehydrated with 20 minute washes at 30%, 40%, 50%, 75%, 80%, 90%, 95%, and 100% ethanol. We used a Tousimis 815 critical point drying system with 100% ethanol. The filters were then placed into a Cressington 108 sputtercoater for 45 seconds and imaged on the JSM-7001F-LV scanning electron microscope at the University of Southern California Core Center of Excellence in NanoImaging (http://cemma.usc.edu/). LSUCC0664 was grown to 10^6^ cells mL^-1^ and 5 µL of culture was loaded onto a glow discharged 300 mesh carbon filmed grid (EMS:CF300-cu). We removed excess liquid with filter paper after 2 minutes and stained with 2% uranyl acetate (TED Pella Cat: 19481) for 1min. The samples were imaged with a JEM-1400 transmission electron microscope at Louisiana State University Shared Instrumentation Facility (https://www.lsu.edu/sif/). We estimated cell volumes using Pappus’ centroid theorem (**Supplemental Text**).

## Results

### New isolate genome characteristics

During the course of previous large-scale culturing experiments, we isolated multiple strains of SAR11 IIIa from the northern Gulf of Mexico (23, 31). We chose three of these isolates (LSUCC0261, LSUCC0664, and LSUCC0723) for further genomic investigation based on their distribution across the 16S rRNA gene tree within SAR11 IIIa (23). Genome sequencing and assembly resulted in a single circular contig for each isolate genome. Characteristically of other SAR11 genomes, our isolate genomes are small (1.17-1.27 Mbp), with low GC content (29-30%), and high coding density (96%) (**Table 1, Fig. S1**).

**Table 1:** Genome statistics of new IIIa isolates compared to other SAR11 genomes. Genome size estimates were calculated by multiplying the assembly size by the inverse of the estimated completion from CheckM (33).

| Genome | LSUCC0664 | LSUCC0723 | LSUCC0261 | Other IIIa | Other SAR11 |
| --- | --- | --- | --- | --- | --- |
| Subclade | IIIa.1 | IIIa.1 | IIIa.3 | IIIa | I,II,LD12 |
| Contigs in Assembly | 1 | 1 | 1 | 1-122 | 1-288 |
| Completion(%)* | 100 | 100 | 100 | 52.38-99.78 | 50.94-100 |
| Est. Contamination(%) | 0 | 0 | 0 | 0-5.95 | 0-4.67 |
| GC(%) | 30 | 29 | 30 | 28-32 | 28-36 |
| Genome Size (Mbp)** | 1.17 | 1.2 | 1.27 | 0.89-1.52 | 0.94-1.75 |
| Coding density (%) | 96 | 96 | 96 | 80-97 | 92-97 |
| Predicted genes | 1256 | 1309 | 1330 | 658-1894 | 654-1788 |

\*Completion criteria of >80% for subclade I/II genomes from GTDB-Tk
\*\*Estimated for genomes that were not closed

### Phylogenomics, taxonomy, and genome trends

Phylogenomics of 471 SAR11 genomes resolved our isolates as novel members of subclade IIIa (**Fig. S2**), and reproduced the three previously observed IIIa subgroups, delineated as IIIa.1, IIIa.2, and IIIa.3 (**Fig. 1A**). While a similar nomenclature was recently proposed (30), we have re-classified the subgroups using results from more genomes, amino acid identity (AAI), and 16S rRNA gene identity (**Fig. 1B**). Both 16S rRNA gene and AAI identities show that IIIa.1 is more similar to IIIa.2 than IIIa.3 (**Fig. 1B**). The lowest 16S rRNA gene identity within IIIa is 92.1% (**Table S1**). Genomes within a subgroup have values of at least 73% AAI to each other with a dropoff of at least 10% AAI between subgroups, which also indicates each subgroup represents genus level classification using AAI (39) (**Fig. 1A-B, Table S1**). Not all of the genomes within IIIa contained a 16S rRNA gene sequence, but those that did shared > 97% 16S rRNA sequence identity within a subgroup. This is near the ∼98% sequence identity metric for species (40). We therefore propose that IIIa represents a taxonomic Family consisting of three genera.

**Figure 1:**
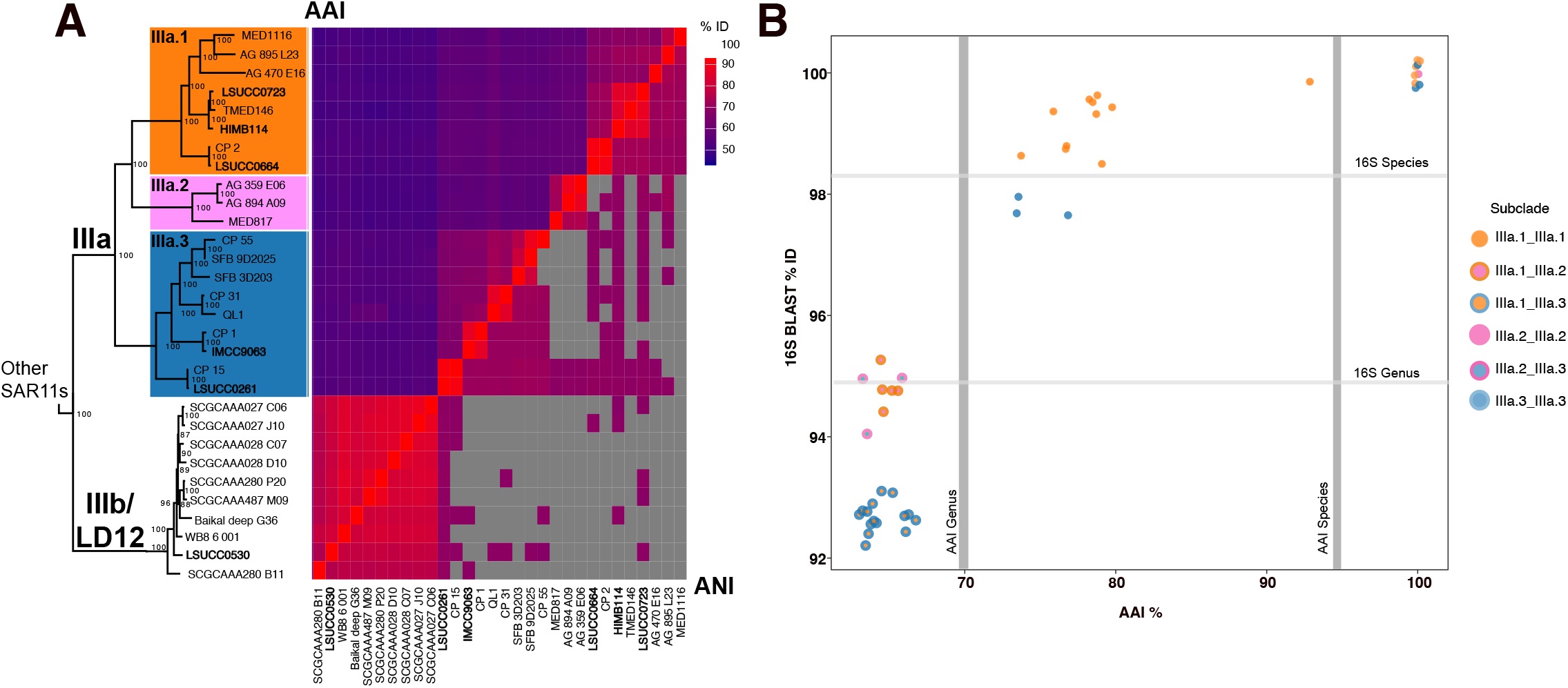
Subclade structure and genome similarity. **A**) Phylogeny and ANI/AAI pairwise comparison of SAR11 IIIa and IIIb. The phylogeny is a subset of the phylogenomic tree found in **Supplemental Figure 1**. Node values are indicators of bootstrap support (n=1000). **B**) 16S rRNA gene BLAST identity vs AAI. Gray bars indicate the species and genera definitions using AAI (78) and 16S (40) where noted.

### Ecological distribution

We removed two non-IIIa MAGs from the SFB that contained contamination > 5% (highlighted in **Table S1**) and recruited reads from 1059 aquatic metagenomes spanning salinities of 0.07-40.2 to 469 SAR11 genomes to evaluate each genome’s relative global distribution across marine and estuarine systems (**Table S1**). We categorized salinity following the Venice system (< 0.5 fresh, 0.5-4.9 oligohaline, 5-17.9 mesohaline, 18-29.9 polyhaline, 30-39.9 euhaline, > 40 hyperhaline) (41, 42) and summed the RPKM values by subclade within a salinity category for each metagenomic sample. Subclade IIIa overall had a wide ecological distribution with habitat specialization by subgroup (**Fig. 2A-B**). IIIa.1 was primarily a polyhaline clade with limited recruitment to sites with salinities < 18. IIIa.2 were euhaline-adapted with the lowest relative abundances of IIIa. IIIa.3 was the most abundant IIIa subgroup in salinities < 30 and appeared primarily adapted for meso/oligohaline environments **Fig. 2B**. Genomes CP31, CP15, LSUCC0261, and QL1 dominated the read recruitment in mesohaline waters and LSUCC0261 was the most abundant isolate genome (**Fig. 2A**), contrasting with the previous use of IMCC9063 and HIMB114 as representatives of the subclade in metagenomic recruitment datasets (22).

**Figure 2:**
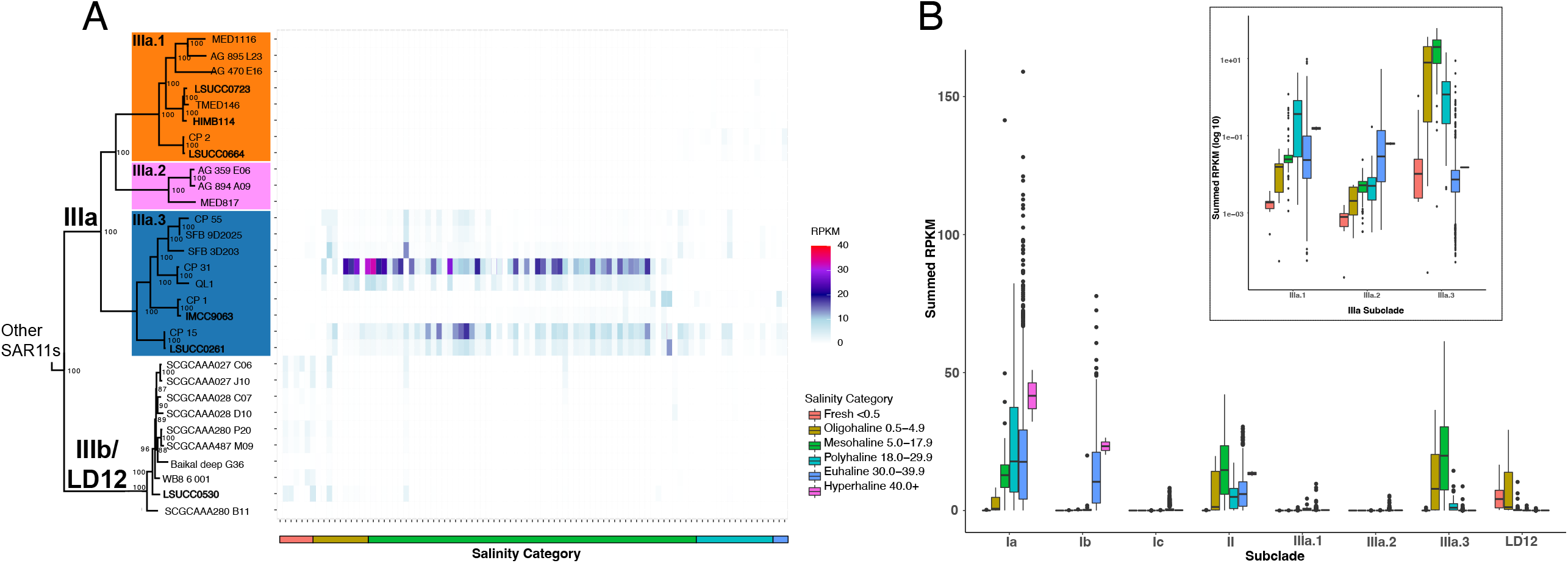
Distribution of subclade IIIa and LD12 in metagenomic datasets. **A)** Metagenomic recruitment to IIIa and IIIb/LD12 genomes at sites with salinities ≤ 32. Tiles represent a metagenomic sample that are arranged by increasing salinity on the x-axis. Colors on each tile represent the Reads Per Kilobase (of genome) per Million (of recruited read base pairs) (RPKM)values at the site. Colors on the x-axis indicate the category of salinity the sample belongs to classified by the Venice system(42). **B)** Boxplot of RPKM values summed by subclade for each metagenomic sample grouped by subclade and colored by salinity category. The insert displays log transformed summed RPKM values for subclade IIIa.

### Genomic content of SAR11 IIIa compared to other SAR11

We conducted a pangenomic analysis of all 471 SAR11 genomes to define genome content similarities and differences within IIIa and between IIIa and other SAR11 with the goals of 1) quantifying differences in metabolic potential, and 2) linking genomic variation to different ecological distributions. Our closed isolate genomes and expanded taxon selection within IIIa allowed us to define whether the previously reported genomic content from IMCC9063 and HIMB114 constituted unique or defining traits of their respective subclades. Although SAR11 potentially contains ten subclades (3) or more (30), for our analysis we condensed these into the broad subclades I, II, and LD12, and excluded subclade V since its membership in SAR11 is controversial (43–47). **Fig. 3** summarizes the genomic differences among SAR11 highlighted below and the complete set of orthologous clusters is in **Table S1**.

**Figure 3:**
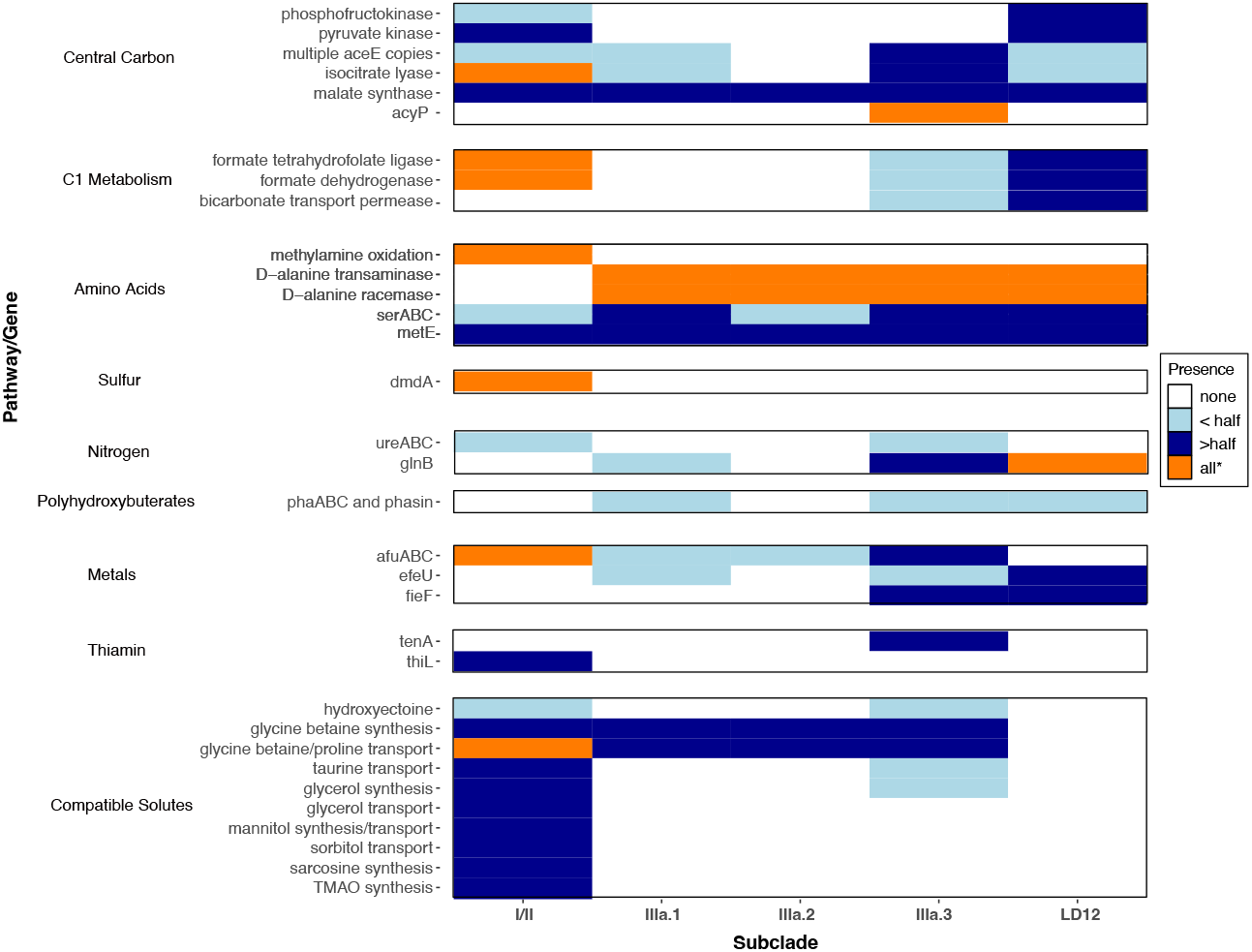
Highlighted comparative gene content in SAR11. Pathways or genes mentioned in text as being differential between subclades are arranged in order of their appearance. Colors indicate the proportion of genomes in a subclade in which the gene/pathway is present in. The asterisk indicates the “all” classification allows for the gene to be missing in limited MAGs or SAGs in subclades I/II since the number of taxa belonging to this group is so large.

#### Central carbo

IIIa had predicted genes for the pentose phosphate pathway, TCA cycle, and glucose 6-phosphate isomerase like subclades I, II, and LD12. IIIa was missing the EMP glycolysis marker gene, phosphofructokinase, that subclades II and LD12 possessed. IIIa was also missing the pyruvate kinase commonly found in LD12 and MAGs and SAGs within subclades I and II. IIIa contained pyruvate dehydrogenase (*aceEF*) like subclades I, II, and LD12. Eight genomes within IIIa contained at least two copies of *aceE*, with QL1 containing 5 copies. Isocitrate lyase is the first enzyme in the glyoxylate shunt that cleaves isocitrate to glyoxylate and succinate. The glyoxylate shunt was not conserved in IIIa (**Fig. 3**), as only 2/8 genomes within IIIa.1 and 5/9 genomes in IIIa.3 contained isocitrate lyase, including LSUCC0664 (IIIa.1) and LSUCC0261 (IIIa.3). However, the closed isolate genome of LSUCC0723 (IIIa.1) did not contain a predicted isocitrate lyase, making it the first reported isolate missing this pathway. The second step of the glyoxylate shunt is carried out by malate synthase, which was common in IIIa and all other subclades of SAR11. Subgroup IIIa.3 uniquely contained *acyP* that breaks an acyl phosphate into a phosphate, carboxyl group, and a proton.

#### C1 metabolism

Most IIIa genomes were missing formate-tetrahydrofolate (THF) ligase and formate dehydrogenase for the production of formate and CO_2_ from the THF-linked oxidation pathway, except for CP31 (IIIa.3) which had both (**Fig. 3**). All IIIa genomes lacked the methylamine oxidation genes that were common in I/II SAR11 as previously reported for HIMB114 (8). Two IIIa.3 genomes, CP31 and LSUCC0261, and six LD12 genomes (including the closed Isolate genome LSUCC0530) contained a sodium-dependent bicarbonate transport permease in the SBT protein family. In freshwater and estuarine cyanobacteria, this protein functions as a high affinity bicarbonate transporter that concentrates inorganic carbon within the cell (48). This probable bicarbonate transporter was found only in CP31 and LSUCC0261 within IIIa.3, which were also two of the genomes that heavily recruited estuary metagenomes (**Fig.2**).Though SAR11 is not known to be able to use inorganic carbon for growth, their genomes do contain carbonic anhydrase and anaplerotic enzymes to use inorganic carbon as intermediates in segments of central carbon metabolism (49).

#### Amino Acids

IIIa and LD12 had the D-alanine transaminase and alanine racemase genes to convert alanine to pyruvate, while other SAR11 did not. Twelve of twenty genomes from IIIa, including our three isolate genomes, contained *serABC* for the production of serine and glycine from glycolysis. Isolates in subclade I were notedly missing the complete gene suite and were consequently reliant on external glycine and serine for their cellular requirements (10, 13), but our analysis found this gene suite present in some MAGs and SAGs within I/II and LD12 (**Fig. 3**). IIIa and LD12 also had multiple copies of a *metE*, a B12-independent methionine synthase. Though this gene was present in I/II genomes, members of IIIa.3 and LD12 had up to three copies spanning multiple orthologous gene clusters (**Table S2**).

#### Sulfur

Like all SAR11, IIIa appear dependent on reduced sulfur compounds and contained no complete assimilatory or dissimilatory sulfate reduction pathways (17, 19). I/II SAR11 were predicted to use DMSO and DMSP, but all IIIa genomes, as well as LD12, were missing *dmdA* for the use of DMSP through the demethylation pathway, confirming the previous observation in the isolate genomes IMCC9063 and HIMB114 (50).

#### Nitrogen and urease

All SAR11 were predicted to use ammonia and synthesize glutamate and glutamine, though the pathways in which glutamate was synthesized were variable. Almost half of IIIa and all LD12 members had *glnB*, a part of the P-II nitrogen response system frequently found in Proteobacteria that is missing in other members of SAR11 (12) (**Fig. 3**). The P-II associated *glnD* gene was not found in any genome, so it is unclear what nitrogen response differences, if any, *glnB* can confer for IIIa/LD12. We found a urease gene suite operon, *ureABC* and accessory proteins *ureEFGHJ* in the isolate LSUCC0261 (IIIa.3) genome with the nickel/peptide ABC transporter commonly found in SAR11. Functional urease operons require a nickel cofactor (51), so the presence of the urease and accessory proteins just downstream the ABC transporter indicated a likely functional gene suite, which we confirmed with growth experiments (below). Thirty-six MAGs from subclade I also contained the urease gene suite (**Table S2**). Urease in SAR11 was first reported in the Eastern Tropical North Pacific oxygen deficient zone where up to 10% of SAR11 were reported to contain the genes (52). Ours is the first reported SAR11 isolate to contain urease and the only extant member of IIIa or LD12 with these genes.

#### Polyhydroxyalkanoates

We found 8/20 genomes within IIIa.1/IIIa.3 and 3/10 genomes in LD12 contained *phaABC* and an associated phasin protein for the predicted production and use of polyhydroxybutyrate (or another polyhydroxyalkanoate) (**Fig. 3**). In other organisms, *phaABC* and phasin proteins allow cells to store carbon intracellularly when carbon is high but another essential component of growth such as nitrogen, phosphorous, magnesium, or oxygen is limiting/unbalanced (53). These granules also have been noted to protect cells from stressors such as temperature, reactive oxygen species, osmotic shock, oxidative stress, or UV damage (54). These genes have been reported in limited IIIa.1 genomes previously (55, 56), but we extend this observation to additional isolates and confirm storage granule synthesis potential as a widespread phenomenon in the IIIa and LD12 subclades. Furthermore, this potential phenotype contrasts with the concept of oceanic SAR11 cells storing phosphate in an extracellular buffer (57). The selection pressure for this gene suite requires further investigation given the broad range of functions for these compounds and the generally high nutrient load of coastal and brackish waters where IIIa and LD12 predominate.

#### Metals

The Fe^3+^ ABC transporter common in subclade I/II SAR11 was found throughout IIIa. Two IIIa.1, three IIIa.3, and seven LD12 representatives as well as HIMB058 (II) also contained *efeU*, a high affinity ferrous iron (Fe^2+^) transporter, and IIIa.3 and LD12 members contained a ferrous-iron (Fe^2+^) efflux pump *fieF* for iron and zinc removal from cells (58) (**Fig. 3**). Estuarine systems have been noted to contain significant amounts of available Fe^2+^ (59), so these genes indicate a potential iron availability niche of which some these specialized SAR11 can take advantage.

#### Compatible solutes

We found GABA (γ-aminobutyric acid) and ectoine synthesis common throughout the SAR11 subclades, but only IIIa.3 members LSUCC0261, CP15, and CP55 (and four SAGS from other subclades) were predicted to synthesize hydroxyectoine from ectoine (**Fig. 3**). Hydroxyectoine is a broad-spectrum osmoprotective molecule for cells, can protect cells against desiccation, and its production was increased during stationary phase when grown in high salt stress in a minimal media in halophile *Virgibacillus halodenitrificans PDB-F2* (60, 61). Glycine betaine synthesis was present in I/II/IIIa and not LD12. The glycine betaine/proline transporter was present throughout IIIa, but IIIa.3 representatives LSUCC0261, CP15, and QL1 were the only members that contain all the subunits, including the ATP binding subunit. This transporter was missing completely in LD12 (19). IIIa.3 members LSUCC0261 and CP15 were the only members of IIIa that could transport taurine like subclades I/II. Glycerol synthesis and transport was present in the I/II subclades, but only two IIIa.3 genomes were predicted to synthesize glycerol from glycerate and no IIIa had genes to transport glycerol. IIIa was also missing mannitol synthesis/transport, sorbitol transport, sarcosine synthesis, and TMAO synthesis though these systems are found in other I/II SAR11. These findings show IIIa contained intermediate numbers of compatible solute genes in between those of I/II and LD12 (**Fig. 3**). IIIa.3 contained the most compatible solute genes within IIIa.

#### Vitamins/cofactors and other genomic features

Six IIIa.3 genomes (including the isolates IMCC9063 and LSUCC0261) and three SAGs outside of IIIa contained *tenA* that should allow the cells to use AmMP rather than HMP as a source of thiamin precursor unlike other SAR11 (14), This distinction is interesting because we also verified that the previously reported loss of the *thiL* gene (14) to phosphorylate thiamin monophosphate to the biologically available thiamin diphosphate (TPP) was conserved throughout subclade IIIa. Thus, although IIIa may exhibit some niched differentiation from Ia via import of a different thiamin precursor, how IIIa produces TPP for use in the cell is unresolved (**Supplemental Text**). Like other SAR11, IIIa had proteorhodopsin–IIIa.1 was a mixture of green and blue (amino acid L/Q at position 105, respectively), IIIa.2 has blue, IIIa.3 has green(**Fig. S3, Supplemental Text**). These spectral tunings correspond to the ecological distribution and source of the genomes with genomes originating from estuarine systems with mesohaline/polyhaline distributions having green. HIMB114, CP1, and AG_894_A09 contained two copies of proteorhodopsin belonging to two orthologous clusters (**Table S2**)– the implications of which are currently unclear and require further study. Isolates LSUCC0723, LSUCC0664, and LSUCC0261 contained no identifiable bacteriophage signatures according to Virsorter (**Table S1**).

### Salinity and temperature growth ranges

We tested the salinity tolerances of two isolates within IIIa, LSUCC0664 (IIIa.1) and LSUCC0261 (IIIa.3) to contextualize the ecological data reported above and understand whether the distribution in ecological data represents the physiological capabilities of the organisms. LSUCC0664 (IIIa.1) grew at salinities of 5.8-34.8 and LSUCC0261 (IIIa.3) grew at salinities of 1.5-34.8, both with an optimum of 11.6. Though the two isolates have an overlapping salinity growth range, LSUCC0261 (IIIa.3) grew faster than LSUCC0664 (IIIa.1) at all salinities except for 23.3 and 34.8, and notably could grow at lower salinities than LSUCC0664 (**Fig. 4A**). These data indicate the IIIa subgroups are euryhaline, in distinct contrast with the sister clade LD12 (19). We also tested isolate LSUCC0261 (IIIa.3) for its temperature range/optimum. It could grow at temperatures of 12-35’C with its optimum of 30’C indicating a preference for warmer waters (**Fig.4B**). While rates of growth between 30-35’C were similar, LSUCC0261 grew to a higher cell density in 30’C (**Figs. 4, S3**).

**Figure 4:**
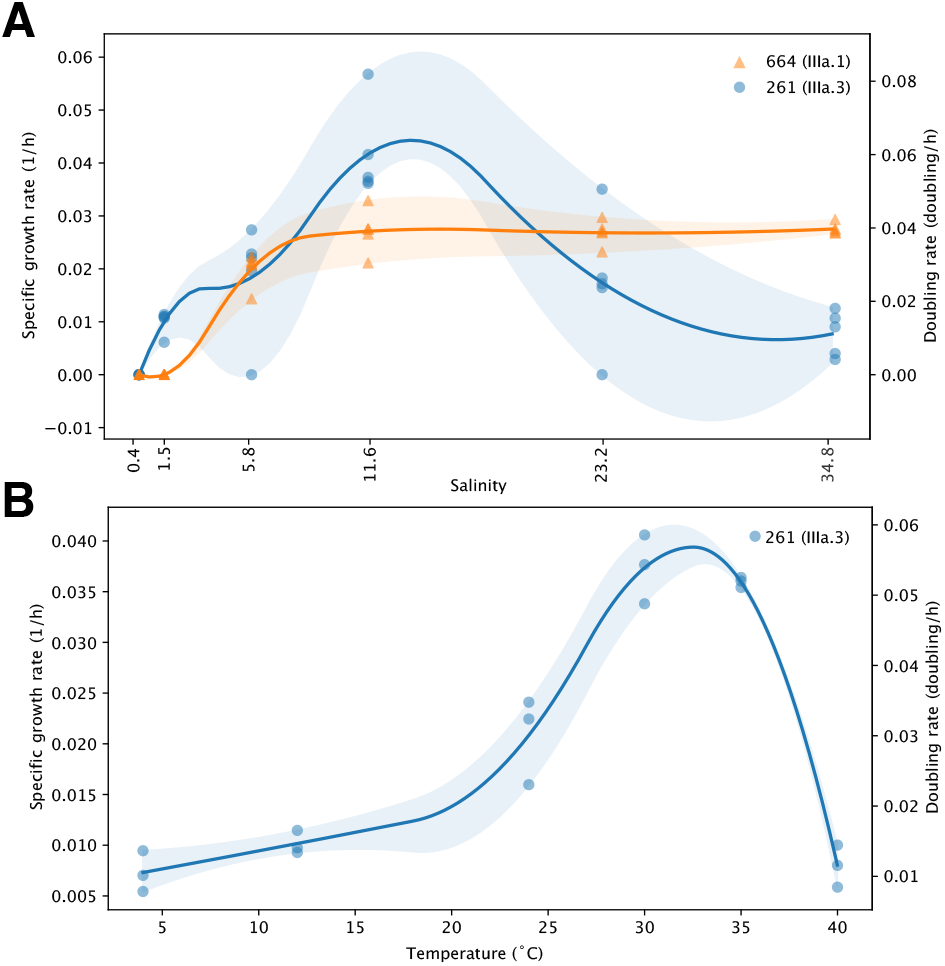
Physiology experiments. **A**) Growth rates and doubling times of LSUCC0664 (IIIa.1) in orange and LSUCC0723 (IIIa.3) in blue in media of varying salinities. **B**) Growth rate and doubling times of LSUCC0261 (IIIa.1) in JW2 medium grown at varying temperatures.

### Minimal C, N, S requirements

We grew LSUCC0261 (IIIa.3) in minimal artificial seawater media to test the isolate’s ability to utilize individual carbon, nitrogen, and sulfur sources with a variety of substrate combinations (**Fig. S5-S6, Table S1**). We tested pyruvate, citrate, ribose, acetate, succinate, and α-ketoglutaric acid as C sources; urea and ammonia as N sources, and cysteine, and methionine as S sources. Oxaloacetic acid, taurine, dextrose, sulfate, DMSO, and DMSP did not support growth. These results are in line with what was predicted by genomics except for oxaloacetic acid which should have been usable as a carbon source due to the presence of *maeB* and its use in isolate HTCC1062 (10). Also in contrast to our study, HTCC1062 was able to use taurine but not acetate as replacements for pyruvate (10) indicating multiple physiological differences between the two isolates.

### Electron Microscopy

Scanning electron microscopy for LSUCC0261 and transmission electron microscopy for LSUCC0664 showed that both cells were curved rods like that of other SAR11 and able to pass through the pores of a 0.1μm laser etched filter (**Fig. 5A-B**). We estimated the cells at 100-300 nm thick for LSUCC0261 and 150 - 240 nm thick for LSUCC0664 (**Fig. S7K**), 0.2 - 1 μm long for LSUCC0261 and 0.4 - 1.5 μm long for LSUCC0664 (**Fig. S7L**), with volumes between 0.01 - 0.05 μm^3^ for LSUCC0261 and 0.015 - 0.04 μm^3^ for LSUCC0664 (**Fig. S7M**). These values are in line with other estimates of SAR11 (62), thus confirming conserved morphology over large evolutionary distances in the Pelagibacterales

**Figure 5:**
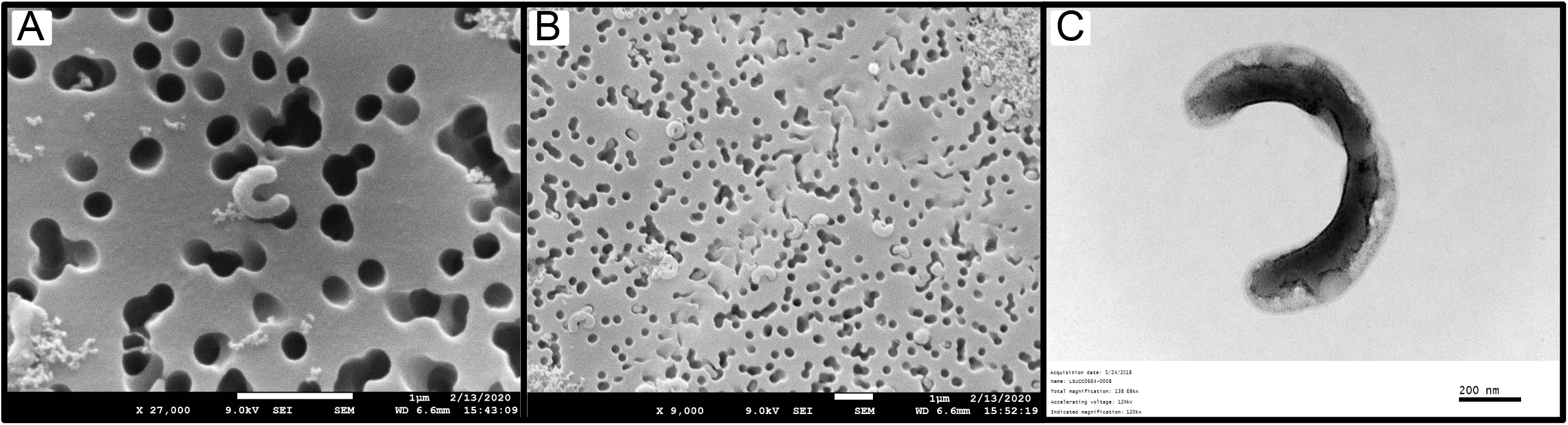
Electron microscopy. **A)** Scanning electron microscopy image of a single LSUCC0261 cell. **B)** Scanning electron microscopy image of many LSUCC0261 cells and cellular debris. **C)** Transmission electron microscopy image of a single LSUCC0664 cell likely mid-division.

## Discussion

This study is the first to systematically focus on SAR11 subclade IIIa and constitutes the most current pangenomic study of high-quality publicly available SAR11 genomes and their phylogenetic relationships. We have contributed multiple new pure cultures and their complete genomes, as well as high quality IIIa MAGs. Previous reports of IIIa genomic content have primarily focused on exceptions to the metabolism of other SAR11 subclades. With our expanded genome selection, we determined whether these findings were conserved features across IIIa or unique to individual isolates. Our study establishes glycine and serine prototrophy; loss of DMSO, DMSP, and much of C1 metabolism; presences of *phaABC* genes; loss of *thiL*; and a mosaic distribution of the glyoxylate shunt as conserved genomic traits within IIIa.

We furthermore confirmed several of these genomic predictions via growth physiology. The isolation of LSUCC0261, LSUCC0664, and LSUCC0723 taxa tested serine and glycine prototrophy because LSUCC0261 was isolated in JW2 medium that does not contain glycine or serine, and LSUCC0664 and LSUCC0723 were isolated in an another medium, MWH2, that did not contain glycine or serine either but did have glycine betaine. HTCC1062 could oxidize glycine betaine as a replacement glycine source (10), but LSUCC0664 and LSUCC0723 do not have the genes to convert glycine betaine to glycine. Thus, the cultivation and propagation of these isolates in our media confirms glycine and serine prototrophy in IIIa. Furthermore, LSUCC0261 did not require glycine or serine in minimal medium experiments (**Fig. S4**) and could not use the reduced sulfur compounds DMSP and DMSO like other SAR11 (28).

This study is the first reported growth of a SAR11 isolate using urea as a sole nitrogen source. Uptake of labeled urea by SAR11 has been observed *in situ* and the urease can be common in OMZ SAR11 (52). While we only observed the urease gene suite in one IIIa genome (LSUCC0261), these SAR11 urease genes were found throughout San Francisco Bay water column metagenomes (**Fig. S8-S9**), suggesting that this metabolism is important for estuarine SAR11. Future work will be needed to: determine whether LSUCC0261 uses urea as a source of nitrogen, carbon, or both; explore the frequency of urease in coastal populations; and identify the circumstances by which urease offers a competitive advantage in SAR11.

Far from being a monolithic subclade with universal features, we propose that subclade IIIa represents a Family within the Order Pelagibacterales and that the subgroups are equivalent to genera defined by both 16S rRNA gene identity and AAI (**Fig. 1**)(39, 40). The genera also had unique spatio-temporal distributions (**Fig. 2B**), which aligns with our understanding of the historical delineation of different SAR11 ecotypes (3, 6, 30, 63). Previous studies defined three phylogenetic branches represented by HIMB114 as a coastal branch (IIIa.1), IMCC9063 (IIIa.3) as a mesohaline branch, and an uncultured oceanic branch between them (3, 22). Our expanded taxon selection and comparison to more than a thousand metagenomes refines our understanding of subclade distribution. While IIIa.3 was the most abundant of the subgroups overall, these organisms preferred slightly lower salinities than IIIa.1, and IIIa.2 was primarily a marine group. Such fine-scale salinity differentiation was supported by physiological data. The IIIa.1 isolate LSUCC0664 could not grow at the lowest salinities possible for LSUCC0261 (IIIa.3) (**Fig. 4**).LSUCC0261 was also best adapted to intermediate salinities, whereas LSUCC0664 grew much better by comparison in higher salinities (**Fig. 4**).

There is important metabolic diversity between the subgroups within IIIa, with IIIa.3 being the most distinct. Several metabolic traits were unique to IIIa.3 or shared only with the freshwater LD12 clade. In addition to the ability to transport Fe^3+^ via ABC transport as other SAR11, IIIa can use a high affinity ferrous iron (Fe^2+^) transporter and IIIa.3/LD12 can pump Fe^2+^ and zinc from cells (58). IIIa.3 contained *acyP* that cleaves acyl-phosphate into a phosphate and carboxylate which may serve as a parallel evolutionary tactic to scavenge phosphate similarly to the methyl phosphonate cleavage in Ia genomes like HTCC7211 (64) or could act simply as an additional way to recycle acetate for the cell’s central carbon metabolism. IIIa.3 has the potential for AmMP to fulfil thiamin requirements instead of being reliant on HMP like most other SAR11(14) due to the presence of *tenA*. In a recent survey of thiamin-related compound concentrations in the North Atlantic, AmMP was found in similar but higher concentrations than HMP at multiple marine stations (65). This represents a crucial niche-differentiating step for IIIa.3 from other SAR11, including the sister groups IIIa.1 and IIIa.2 that are likely reliant on HMP (14). Subclade IIIa’s conserved deletion of *thiL*, which converts thiamin monophosphate (TP) to the biologically usable thiamin diphosphate (TPP), remains inexplicable as it appears that these organisms still require thiamin diphosphate. For example, eight genomes spanning the three subgroups within IIIa have multiple gene copies of the *aceE* E1 component of pyruvate dehydrogenase and QL1 has five copies. This is notable because gene duplications in SAR11 are limited (8), and also because *aceE* needs thiamin diphosphosphate as a cofactor to combine thiamin diphosphate and pyruvate to make acetyl-CoA (66). It is thus likely that a currently unannotated gene can complete this final conversion. One possible candidate is an adenylate kinase found in all SAR11 that can convert thiamin diphosphate to thiamin triphosphate (67). Combined, these notable metabolic shifts in IIIa.3 probably allow for the subclade to exploit environmental resources that other SAR11 are unable to use and contribute to the ecological success of the group relative to the other groups in IIIa.

Truly estuarine-adapted taxa are believed to be rare compared to marine and freshwater versions (68). Prior research from river outlets debated whether estuarine-adapted lineages could truly exist or whether the community members in estuarine zones are simply a mixture of freshwater and marine communities because the short residence times of estuarine water make an established community unlikely (69). However, a true brackish community in the Baltic Sea between salinities of 5-8 was distinct from fresh and salty community members (70). The physiology, ecological distribution, gene content, and sister position of IIIa to LD12 all support the concept of an estuarine origin of the last common ancestor for IIIa/LD12. Subsequently, one subgroup of IIIa remained truly estuarine-adapted (IIIa.3), whereas the other subgroups diversified into increasingly higher salinity niches over time (IIIa.1 and IIIa.2). While marine to freshwater transitions are rare (71), freshwater to marine transitions are perhaps even more rare. Bacteria such as the Methylophilaceae have recently been documented to have freshwater origins for marine relatives (72) and some diatoms such as the Thalassiosirales have extensive marine to freshwater transitions followed by subsequent marine transitions(73). While IIIa appears to be a transitionary clade diversifying from estuarine waters back to marine systems, more genomes and further research into physiology and biogeography are needed to improve our understanding of the evolutionary origins and trajectory of this group.

More generally, subclade IIIa represents an intermediate group in the SAR11 evolutionary transition from marine to fresh water. These organisms inhabit a wide range of salinities but are brackish water specialists and share a most recent common ancestor with the exclusively fresh and low-brackish water subclade LD12. The last common ancestor of all SAR11 is believed to be a streamlined, marine organism (74), and we currently hypothesize that a key evolutionary step that allowed the colonization of fresh water occurred through the loss of osmolyte transport genes (for glycine-betaine, proline, ectoine, and hydroxyectoine) in the LD12 branch (19). The tradeoff for this gene loss was that LD12 was prevented from reinhabiting salty waters (19). We can use the knowledge of subclade IIIa gained from this study to speculate on this evolutionary transition further. The two isolates, LSUCC0261 and LSUCC0664, have a euryhaline growth range. While this is noteworthy by itself, it is perhaps more important that LSUCC0261 cannot grow in the lowest salinity media, i.e., fresh water. What prevents this growth at the freshest salinities remains an important question. Key features of SAR11 are small, streamlined genomes that have a comparative dearth of regulatory capability (9) and a high number of constitutively expressed genes (75). It is possible that the constitutive expression of the very same osmolyte transport genes that were lost in LD12 prevents IIIa cells from reducing their ionic strength sufficiently to inhabit fresh water.

The path of SAR11 evolution from the marine clades to LD12 would follow thusly: specialization for brackish water habitats first occurred at the branch between all SAR11 and the last common ancestor of IIIa/LD12. These organisms were distributed very near freshwater environments but could not permanently colonize them due to constitutive gene expression of osmolyte transport genes that prevented sufficient reduction in intracellular salinity. Loss of these transport genes then allowed the LD12 group to disperse into fresh water, but the absence of these genes prevented rapid equilibration to higher salinities in the event of re-dispersal back to marine waters, isolating LD12 as a freshwater group (19). We are currently investigating this hypothesis with isolates from IIIa and LD12. And while the evolutionary trajectory for LD12 may pass through the common ancestor of IIIa and LD12 as outlined above, there is accumulating evidence that the ostensibly exclusively marine SAR11 groups may also colonize freshwater environments either sporadically, at very low abundances, or both (76, 77)

Overall, this study represents the most complete analysis of SAR11 IIIa thus far and is a necessary steppingstone in the understanding of SAR11 IIIa, its role in estuarine systems, and its intermediate place in the evolution of SAR11 from marine to freshwater environments. Future work on IIIa is needed to contextualize functions of noted gene losses and gains, the mode in which IIIa interacts with thiamin derivatives, and the extent at which IIIa members interact with nutrient dynamics in estuaries including urea and production of polyhydroxyalkanoates.

## Data Availability Statement

Assembled isolate genomes for LSUCC0261, LSUCC0664, and LSUCC0723 are available on IMG under Genome IDs 2728369215, 2770939455, and 2739368061, respectively. Raw isolate genome reads are available on NCBI under accession PRJNA864866. Metagenome assembled genomes from the San Francisco Bay are available on NCBI under BioSample accessions SAMN30106608-SAMN30106615. The accessory datasheets from this publication including the pangenome summary are hosted through FigShare (https://figshare.com/account/home#/projects/144939). Cryostocks of isolates used in this analysis are available upon request.

## Acknowledgements

We would like to thank the Louisiana State University Shared Instrumentation Facility (SIF) and the University of Southern California Center for Electron Microscopy and Microanalysis (CEMMA) for training and availability of electron microscopes to image our isolates. We would also like to thank Dr. Casey Barr for training on the scanning electron microscope and Dr. Ying for her operation of the transmission electron microscope. The authors acknowledge the Center for Advanced Research Computing (CARC) at the University of Southern California for providing computing resources that have contributed to the research results reported within this publication. URL: https://carc.usc.edu, as well as high-performance computing resources provided by Louisiana State University (http://www.hpc.lsu.edu), and the Stanford Research Computing Center for providing computing resources that have contributed to the research results reported within this publication. This work was supported by a Simons Early Career Investigator in Marine Microbial Ecology and Evolution Award, and NSF Biological Oceanography Program grants (OCE-1747681 and OCE-1945279) to J.C.T.

## Figure Captions

**Supplemental Text 1**: Supplemental Methods, Results, and Discussion.

**Supplemental Table 1**: Accessory data used in this publication including: GTDB accessions, CheckM statistics, and estimated genome size for all genomes, table of noted genomic features in text, 16S blast hits of IIIa, ANI and AAI matrix of IIIa, AAI vs BLAST of IIIa, detailed gene searches corresponding to previous publications, table of KO numbers that differ between LSUCC isolate genomes, Anvi’o enriched pfam and KO, Virsorter outputs for isolates, input table for sparse_growth_curve.py to calculate growth rates from salinity and temperature experiments, growth data for minimal media experiment, minimal media setup, metagenomic recruitment RPKM values, and collected metadata for the datasets used in recruitment. Supplemental Table 1 is hosted at: https://doi.org/10.6084/m9.figshare.20415831.

**Supplemental Table 2**: Anvi’o pangenomic summary of 471 SAR11 genomes annotated with the following sources from KEGG and Interproscan: Gene3D, SUPERFAMILY, TIGRFAM, KEGG_Class, KOfam, ProSiteProfiles, Pfam, CDD, Hamap, PANTHER, KEGG_Module, PIRSF, SMART, ProSitePatterns, Coils, MobiDBLite, PRINTS, SFLD. Supplemental Table 2 is hosted at: https://doi.org/10.6084/m9.figshare.20415843.

**Supplemental Table 3:** Cell sizes measurements and estimations in **Fig. S7**. Supplemental Table 3 is hosted at: https://doi.org/10.6084/m9.figshare.20415852.

**Figure S1**: Boxplots of genome characteristics of LSUCC isolates compared to other SAR11.

**Figure S2**: Phylogenomic tree of 471 SAR11 genomes that are a combination of newly-added genomes and publicly available. Node values are indicators of 1000 bootstrap support.

**Figure S3**: Multiple sequence alignment of IIIa proteorhodopsin with key spectral tuning position boxed in red.

**Figure S4:** LSUCC0261 growth at different temperatures. Points indicate the average of three replicates and error bars indicate the standard deviation of cell counts for three replicates measured.

**Figure S5:** LSUCC0261 growth in different minimal media. Points indicate the average of three replicates and error bars indicate the standard deviation of cell counts for three replicates measured.

**Figure S6:** Growth rates of LSUCC0261 grown in different minimal medium combinations.

**Figure S7**: Calculations of cell sizes. For example with the annotation in (**G**): we have the same circles covering the entire cell shape. The radii (R = 45px = 88nm, half of the cell thickness) of the identical circles includes the two half-spheres and the curved cylinder, from which we can calculate the volume of the two half-spheres (in total, 4/3πR^3^ = 0.0029 μm^3^). We then connect the centers of the circles. The length of the connection line (l = 633.7 px = 1239 nm) is the length of the curved cylinder. According to Pappus’ centroid theorem, the volume of the curved cylinder is πlR^2^ = 0.0301 μm^3^. We then get the total cell volume as 0.033 μm^3^. We applied this method to all the cells in the images. (**A**) – (**D**) are the scanning electron microscopic images for LSUCC0261. (**E**) and (**F**) are the transmission electron microscopic images for LSUCC0261. (**G**) – (**J**) are the transmission electron microscopic images of LSUCC0664. (**G**) and (**I**) are two identical images, where (**G**) is being annotated as a whole single cell while (**I**) is annotated as two newborn cells since due to the presence of a likely septum. As the result, (**K**) — (**M**) are showing the distributions of cell radius, lengths, and volumes. The data for making the violin plots are in **Table S3**.

**Figure S8**: Phylogeny of the UreC sequences from the San Francisco Bay (SFB) with the LSUCC0261 sequence highlighted.

**Figure S9**: Plot of BLAST hit percent identities between the SFB UreC sequences and the LSUCC0261 UreC organized by samples with increasing salinities.

