## Supplemental text and figures for "Ecophysiology and genomics of the brackish water adapted SAR11 subclade IIIa"

Supplemental text for **Ecophysiology and genomics of the brackish water adapted SAR11 subclade IIIa**

### Supplemental Methods

*Isolation, genome sequencing and assembly (continued)*

LSUCC0261 was isolated using JW2 medium in September 2015 and LSUCC0664 and LSUCC0723 were isolated using MWH2 medium in September 2016, from the Calcasieu Jetties in Cameron, LA (29.760164 -93.340159)(1, 2). We chose these strains for genome sequencing because their location on the 16S rRNA gene phylogenetic tree indicated that they represented two of the existing branches of IIIa (1, 2). For genome sequencing, we grew LSUCC0664 and LSUCC0723 in MWH2 and LSUCC0261 in JW2 media and filtered them through a 25 mm 0.22 µm polycarbonate filter (Millipore, Massachusetts, USA) when cultures reached roughly $10^6$ cells $mL^{-1}$. DNA was extracted with the MoBio PowerWater DNA kit (QIAGEN, Massachusetts, USA) following the manufacturer's protocol and eluted in 50uL of Mili-Q water.

DNA for strain LSUCC0261 was sequenced using an Illumina HiSeq after library preparation as previously reported (3) at the Oklahoma Medical Research Facility. Sequencing produced 6,134,004 paired-end 151bp reads with a 400bp insert size. We selected a subset of 1,000,000 reads for assembly using seqtk as previously reported (3). DNA for strains LSUCC0664 and LSUCC0723 was sent to the Argonne National Laboratory Environmental Sample Preparation and Sequencing Facility for library preparation and sequencing. These Illumina TruSeq libraries were sequenced using and Illumina MiSeq and generating 1,609,453 and 2,314,532 paired-end 251bp reads with a 550bp insert size. We trimmed the resulting sequences with Trimmomatic v0.36 using LEADING:20 TRAILING:20 SLIDINGWINDOW:13:20 MINLEN:40 for LSUCC0261 and LSUCC0723 and LEADING:20 CROP:150 SLIDINGWINDOW:13:20 MINLEN:40 for LSUCC0664 and assembled trimmed reads for all genomes with SPAdes v3.10.1 (4) with --cov-cutoff option set to "auto". The final assembly coverages, calculated via Pilon, were as follows: LSUCC0261- 1112x, LSUCC0664- 491x, LSUCC0723- 699x. Default settings were used for all software during the genome assembly process unless otherwise noted.

*Confirmation of closed genomes*

The LSUCC0261 and LSUCC0723 assemblies resulted in single scaffolds with overlapping ends. We evaluated the strain LSUCC0261 assembly using similar methods as previously reported (1) involving two rounds of Reapr v1.0.18 (2) with one round of SSPACE v3.0 as reported (3) using all HiSeq reads, as well as reapr/SSPACE runs on the rotated scaffold that was manually joined at the ends and broken on the opposite side. This was followed by evaluation of the rotated scaffold with Pilon v1.22 (4) with default settings using the indexed bam files from reapr/SSPACE. Pilon made no corrections. We evaluated the LSUCC0723 assembly using Pilon only, after mapping the reads to the scaffold using BWA (5), and repeated this process with the rotated scaffold as for LSUCC0261, above. Pilon made no corrections on either run.

The LSUCC0664 assembly resulted in a single scaffold with overlapping ends, but while Pilon (completed as described above) returned no corrections on the original scaffold, it found two continuity breaks when we rotated the scaffold at the location where the ends overlapped in the original scaffold (573,860-574,412 and 574,938-575,012). Thus, we designed primers to verify the overlap with the NCBI Primer-BLAST website (https://www.ncbi.nlm.nih.gov/tools/primer-blast/) using the following settings: Forward primer from 573800 to 573900; Reverse primer from 574950 to 575050; PCR product size Min 900 / Max 1300; Tm Min 52 / Opt 54 / Max 55; remainder default. We amplified DNA using the following primers: F-ATAATGTCAAACTTGCACCTG, R-TCTAAGATTTACCCCTGTAGC, which were predicted to set down at bases 573,834-573,854 and 575,003-575,023, respectively, outside of the region of interest. PCR was conducted with the following annealing temperatures: 53, 52.5, 51.7, 50.4, 48.8, 47.6, 46.7, 46 (all ˚C). Multiple bands were observed at each temperature. We selected the heaviest band, purified the gel, and Sanger sequencing of the resultant PCR product produced overlapping reads from positions and 573,932-574,841 from the F primer and 574,018-575,034 from the R primer, confirming that this section of the scaffold was contiguous.

*Taxon selection (continued):*

We used known SAR11 isolate genomes listed in **Table S1** as input for GTDB-Tk v1.5.0 – classify_wf (5) to clearly define the SAR11 clade *de novo* and screened all genomes from the SAR11 node of the GTDB output using the CheckM metadata. We required completeness of at least 50% and contamination of < 5% to define a starting set. We then culled the genome selection using the following criteria: any IIIa genome that was > 50% complete and < 5% contamination was kept to ensure the fullest possible sampling of available genomes, whereas genomes within subclades I and II were maintained only if they had > 80% completion and < 5% contamination because of an overrepresentation of genomes from these subclades in GTDB. We supplemented the LD12 subclade with 5 additional genomes downloaded from IMG (2018) to better match taxon selection in our previous work (3).We then dereplicated the resulting genomes using fastANI (6) with default settings, defining duplicates as those with > 99% average nucleotide identity (ANI), and kept the genome from any matching pair with higher completion and lower contamination as estimated using CheckM v1.0.5 (7). This final genome set was combined with MAGs assembled from the San Francisco Bay (SFB) (see below and REFXX) that were at least 50% complete with < 8% contamination as estimated with CheckM v1.0.5 (7) and our isolate genomes for a total of 471 genomes.

*MAG assembly from the SFB*

MAGs from SFB were generated using the methods described previously (8). Data is deposited in BioProject no. PRJNA819083.

*Phylogenomics*

We processed all genomes through the Anvi'o version 6.2 pipeline (9) as referenced previously (10) to gather single-copy marker genes present in at least half of the genomes. The 70 resulting amino acid sequences were aligned with MUSCLE v3.8.1551 (11), trimmed with trimAl v1.2 (12), concatenated with genestitcher.py, and the final tree was inferred with IQ-TREE (version 1.6.12, default settings) (13). We used FigTree v1.4.4 (http://tree.bio.ed.ac.uk/software/figtree/) for visualization of the final tree.

*Proteorhodopsin tuning identification*

To investigate proteorhodopsin in SAR11 IIIa, we gathered amino acid sequences annotated as rhodopsins from the pangenome summary (**Table S2**), aligned them with MUSCLE v3.8.1551 (11), and visualized the alignment with NCBI's Multiple Sequence Alignment Viewer v.1.21.0 (**Fig. S3**). We compared amino acid position 105 (slightly different in our alignment because of gaps) to identify the spectral tuning of each genome's rhodopsin.

*SFB urease gene tree:*

To build a single-gene phylogeny of the SAR11 urease gene, we ran blastp on the LSUCC261 large subunit of the urease protein (UreC) to the nr database (accessed June 2018), collected the 200 best hits while excluding those labeled as "MULTISPECIES", and added UreC amino acid sequences from the San Francisco Bay metagenomic dataset (8) that were annotated as ureC in IMG and identified using either a KO or pfam function search. The sequences were aligned with MUSCLE v3.8.1551 (11), trimmed with trimAl v1.2 (12) using -automated1, and inferred with IQ-TREE (version 1.6.12, default settings) (13).

*Cell size estimation*

To calculate the volume of the cell, we first separated the cell image into two half spheres and a curved cylinder. The cell volume, therefore, is the sum of the volumes of the curved cylinder and the two half-spheres, where the radii of the half-spheres equal the section of the curved cylinder. We estimated the length of the curved cylinder by drawing a curve connecting the center points of all sections of the cylinder. The volume of the curved cylinder is the area of the section ($\pi r^2$) x length (Pappus' centroid theorem). The volume of the two half-spheres was combined into one as $4/3\pi r^3$. We summed the volumes of the sphere and the curved cylinder for the final estimate of cell volume. The details of the calculations can be seen in **Fig. S6 and Table S3**. If a clear septum is formed according to the image, we will do two separate annotations on the same image, one for the parent cell **(Fig. S7G**), the other one for the two children cells (Fig.S6I). As the result, the sizes of the children cells are the approximate minimum size of the strain and the size of the mother cell is the approximate maximum.

### Supplemental Results and Discussion

*Additional genomic content not included in main text*

Energetics. IIIa genomes contained oxidative phosphorylation genes encoding for an aerobic chemoorganoheterotrophic lifestyle- as in other SAR11 genomes- with a predicted cytochrome c oxidase, F-type ATP synthase, and a proton-pumping NADH dehydrogenase. Five members of IIIa.1 and two members of IIIa.2 including isolates within IIIa.1(HIMB114, LSUCC0664, and LSUCC0723) contained multiple copies of cytochrome c that belonged to two separate orthologous gene clusters.

Membrane Transport. Like all other subclades of SAR11, IIIa contained polar amino acid ABC transport, *ntrXY* genes for nitrogen sensing and assimilation, ammonium transport via *amtB*, the *regAB* redox sensing two-component system, the *envZ* osmoregulation two-component system, C4 dicarboxylate Tripartite ATP-Independent Periplasmic (TRAP) transporter, a phosphate ABC transporter, and the *sec*-dependent pathway.

Minimal media verification of glycine/serine prototrophy.

We report the first verification of SAR11 IIIa glycine and serine prototrophy evidenced by growth in JW2 and minimal media combinations found in **Fig. S5**. Minimal media combinations that support growth included some TCA cycle intermediates that in theory could be used to form glycine from glycolate via the glyoxylate shunt. It was hypothesized for HTCC1062 that the presence of the glyoxylate shunt wouldn't supply sufficient glycine for biomass because the shunt directed the carbon compounds away from biosynthesis and towards energy production due to low glycine concentrations in the cell (14). If TCA cycle intermediates were to produce glycine/serine via the glyoxylate bypass, the glycine concentration in the cell would already need to be substantial through prototrophy. Additionally, LSUCC0723 does not contain the glyoxylate shunt and grew in MWH2 medium, verifying the glycine/serine prototrophy is real rather than supplied via TCA intermediates.

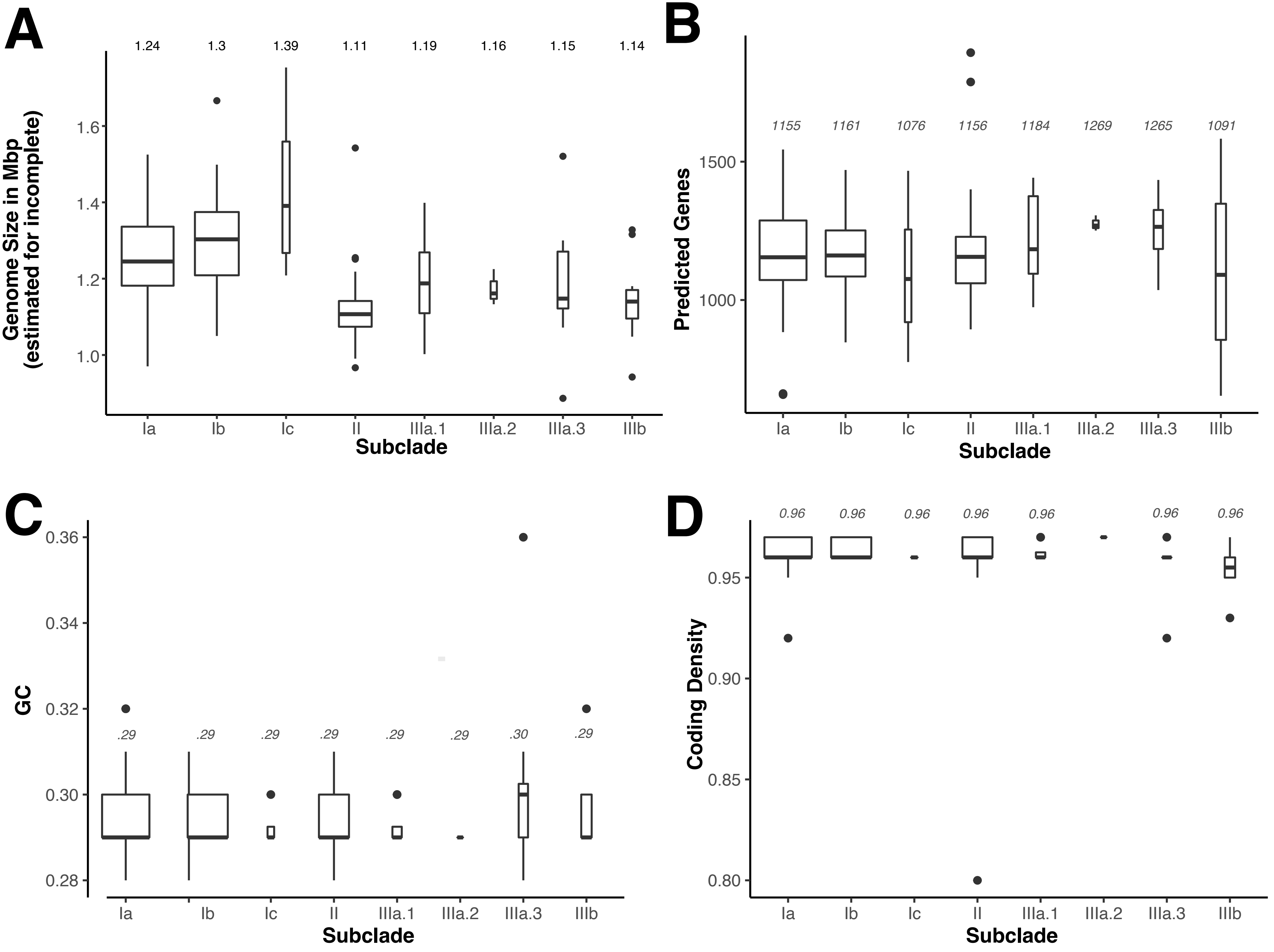


**Figure S1**: Boxplots of genome characteristics of LSUCC isolates compared to other SAR11.

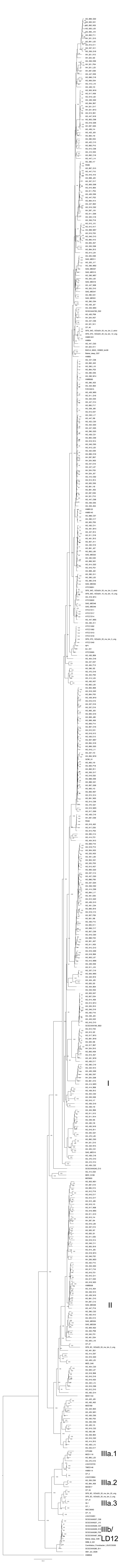


**Figure S2**: Phylogenomic tree of 471 SAR11 genomes that are a combination of newly-added genomes and publicly available. Node values are indicators of 1000 bootstrap support.

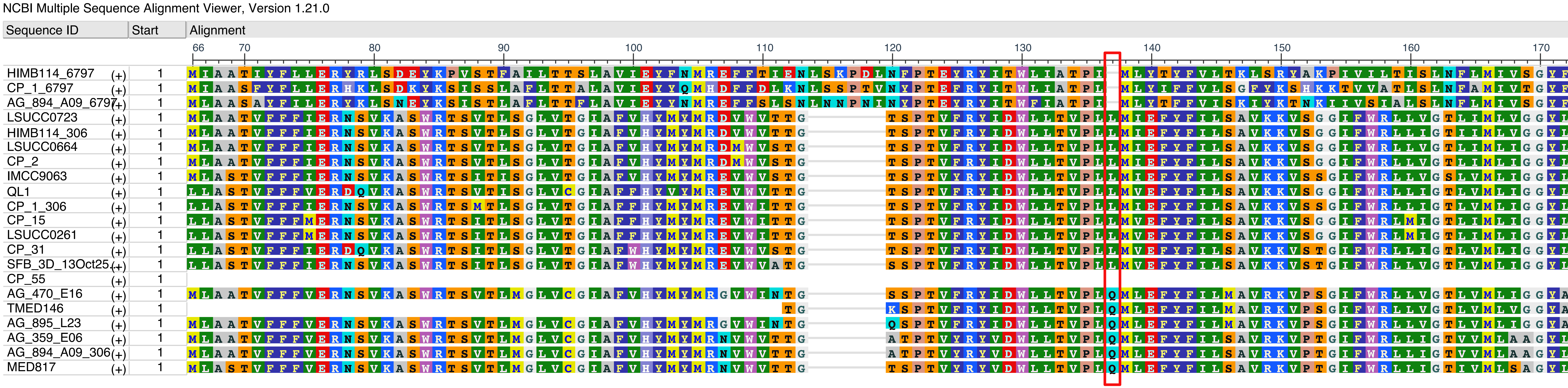


**Figure S3**: Multiple sequence alignment of IIIa proteorhodopsin with key spectral tuning position boxed in red.

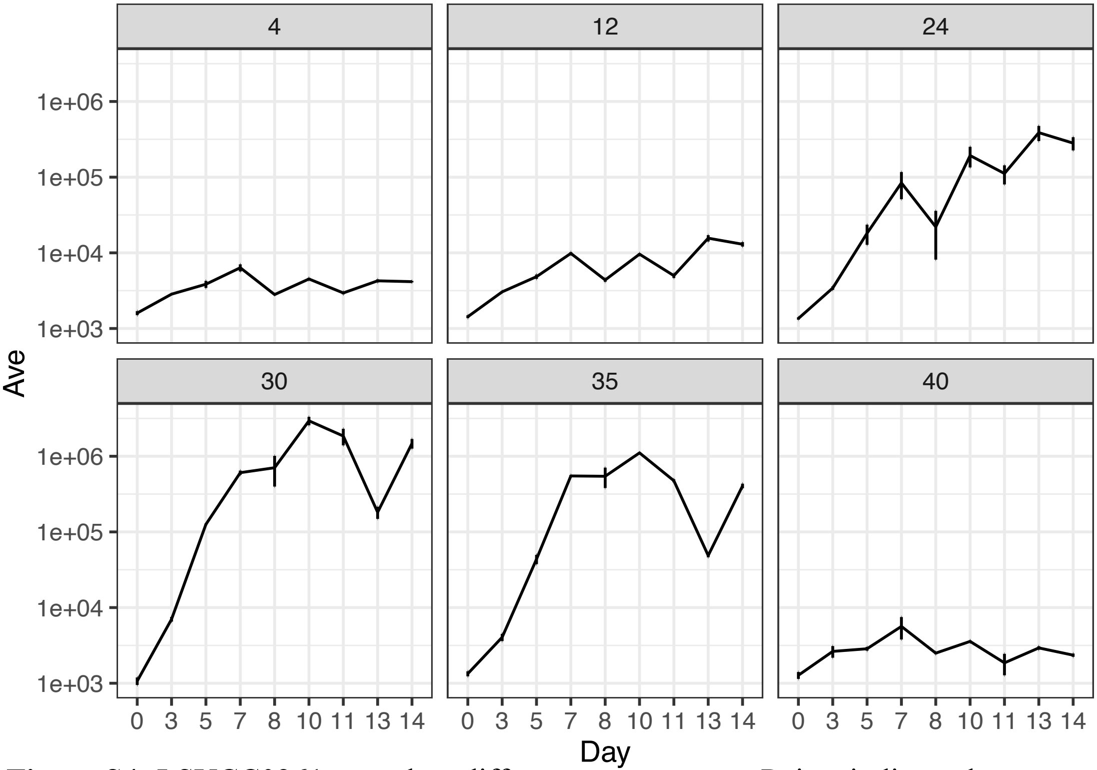


**Figure S4:** LSUCC0261 growth at different temperatures. Points indicate the average of three replicates and error bars indicate the standard deviation of cell counts for three replicates measured.

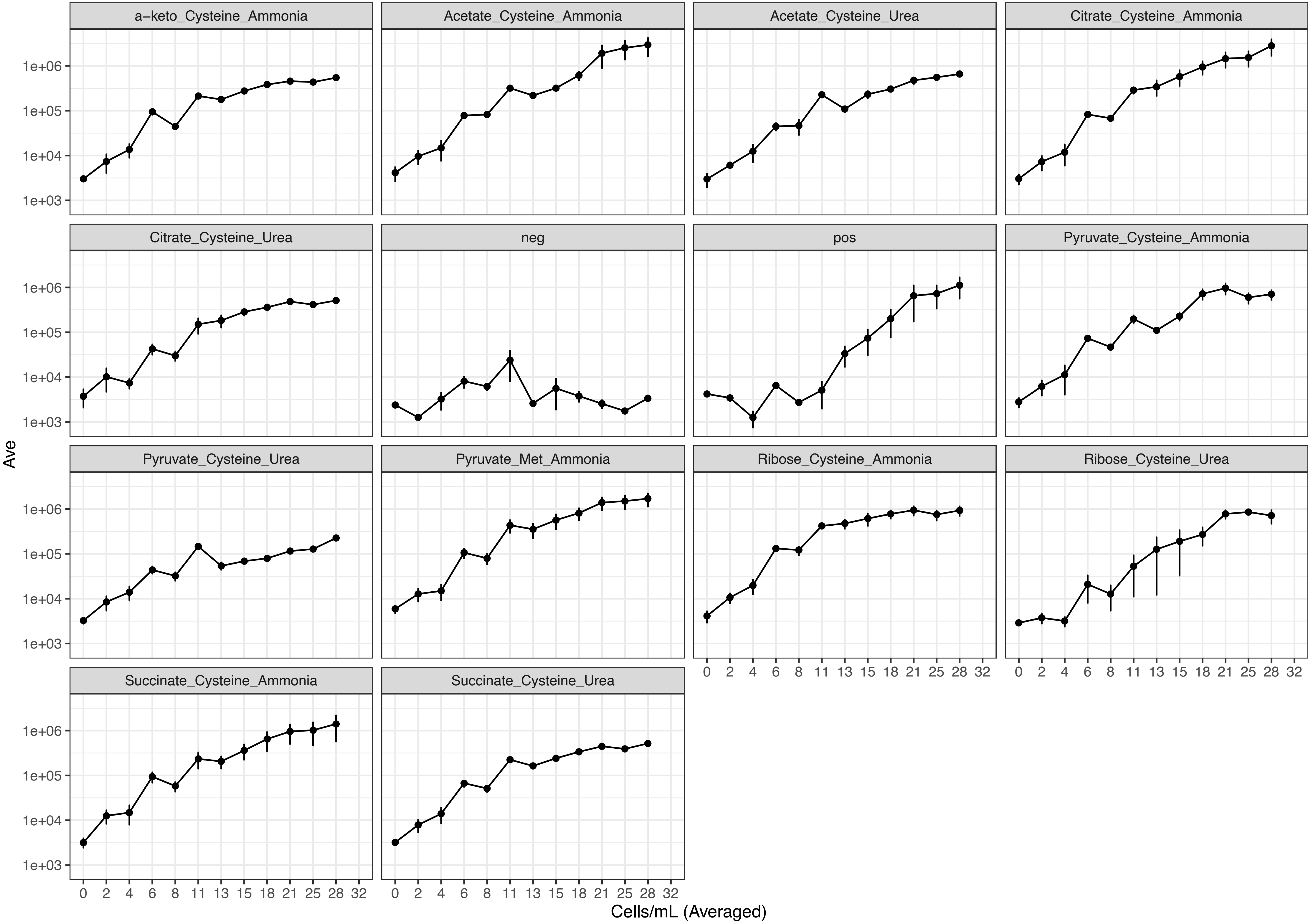


**Figure S5:** LSUCC0261 growth in different minimal media. Points indicate the average of three replicates and error bars indicate the standard deviation of cell counts for three replicates measured.

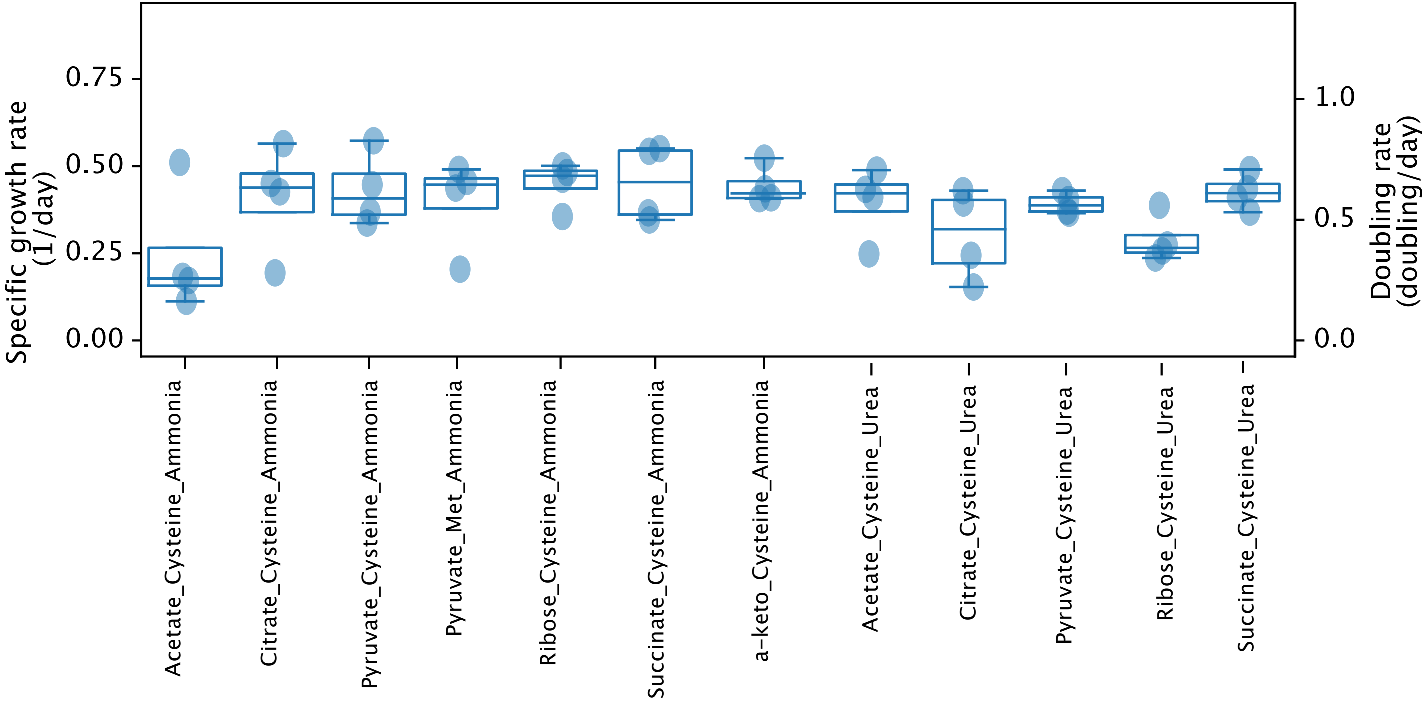


**Figure S6:** Growth rates of LSUCC0261 grown in different minimal medium combinations.

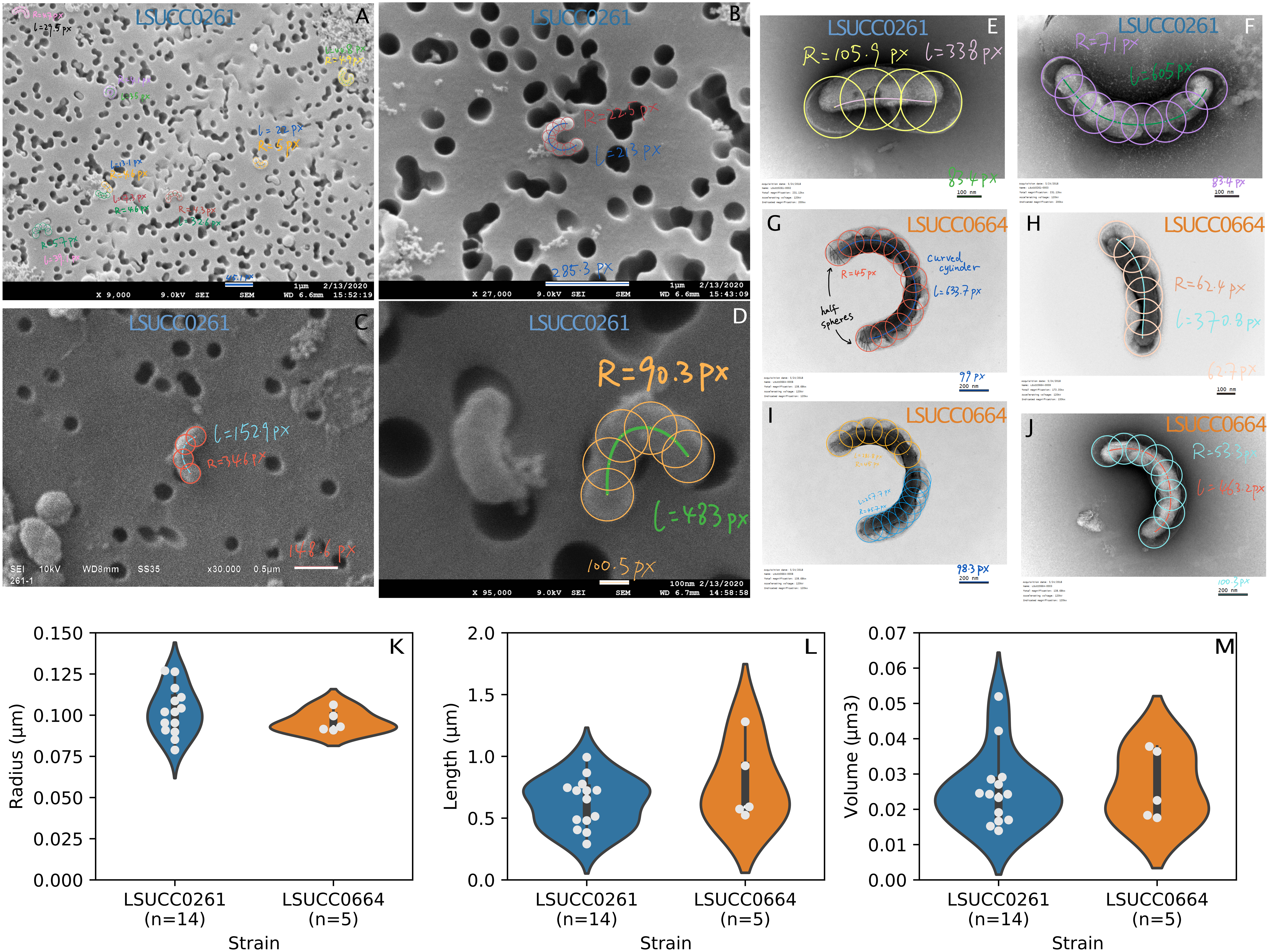


**Figure S7**: Calculations of cell sizes. For example with the annotation in (**G**): we have the same circles covering the entire cell shape. The radii (R = 45px = 88nm, half of the cell thickness) of the identical circles includes the two half-spheres and the curved cylinder, from which we can calculate the volume of the two half-spheres (in total, $4/3\pi R^3$ = 0.0029 μm$^3$). We then connect the centers of the circles. The length of the connection line (l = 633.7 px = 1239 nm) is the length of the curved cylinder. According to Pappus' centroid theorem, the volume of the curved cylinder is $\pi l R^2$ = 0.0301 μm$^3$. We then get the total cell volume as 0.033 μm$^3$. We applied this method to all the cells in the images. (**A**) – (**D**) are the scanning electron microscopic images for LSUCC0261. (**E**) and (**F**) are the transmission electron microscopic images for LSUCC0261. (**G**) – (**J**) are the transmission electron microscopic images of LSUCC0664. (**G**) and (**I**) are two identical images, where (**G**) is being annotated as a whole single cell while (**I**) is annotated as two newborn cells since due to the presence of a likely septum. As the result, (**K**) — (**M**) are showing the distributions of cell radius, lengths, and volumes. The data for making the violin plots are in **Table S3**.

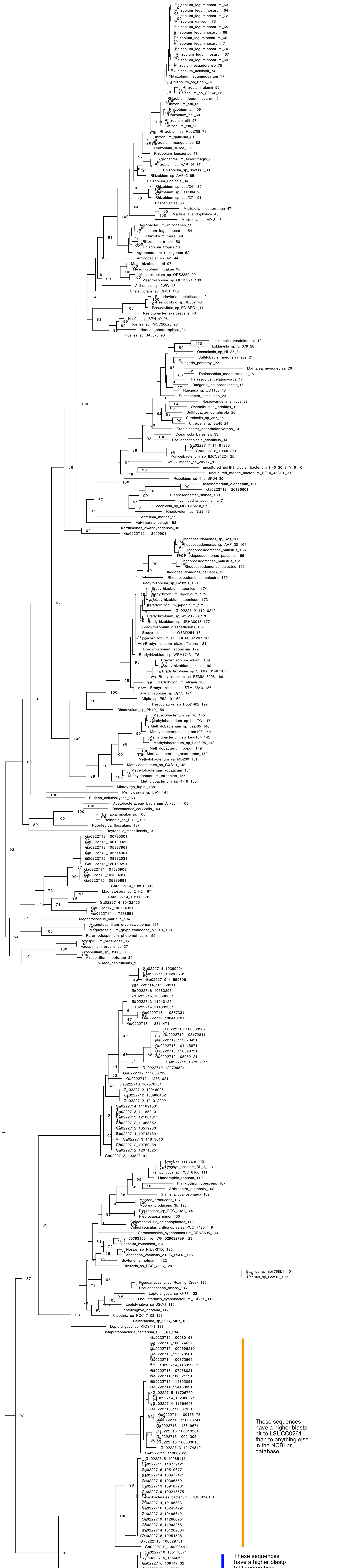


**Figure S8**: Phylogeny of the UreC sequences from the San Francisco Bay (SFB) with the LSUCC0261 sequence highlighted.

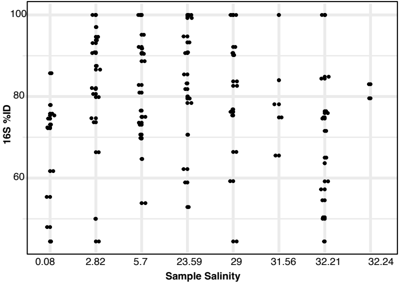


**Figure S9**: Plot of BLAST hit percent identities between the SFB UreC sequences and the LSUCC0261 UreC organized by samples with increasing salinities.
